# The Duplicate Monophyly Criterion: An Empirical Approach to Bootstrapping Distance-Based Structural Phylogenies

**DOI:** 10.64898/2026.03.25.713827

**Authors:** Ashar J. Malik, David B. Ascher

## Abstract

Distance-based structural phylogenies summarise relationships between proteins using pairwise structural similarity scores. Unlike character-based methods, however, they lack a natural analogue of the non-parametric bootstrap. Because distances derived from continuous, high-dimensional folds provide no discrete sites to resample, rigorous support estimation would ideally rely on conformational ensembles from Molecular Dynamics or Monte Carlo simulations, which is computationally prohibitive at scale. A practical alternative is parametric bootstrapping in distance space, but this introduces a calibration problem: without an objective estimate of the signal-to-noise ratio, the perturbation magnitude cannot be chosen in a principled way. Here, we introduce the *Duplicate Monophyly Criterion* (DMC), an empirical strategy for calibrating distance-matrix perturbations using synthetic taxon duplicates as internal controls. We expand the observed distance matrix with virtual copies of each taxon, assign each original–duplicate pair a small baseline distance, and apply a floor-augmented heteroscedastic noise model directly in distance space. The central hypothesis is that loss of duplicate monophyly (i.e., failure of taxon–duplicate pairs to form two-tip cherries) marks a regime in which perturbations overwhelm intrinsic phylogenetic signal, thereby defining a conservative operating range for stability-based (bootstrap-like) support estimation. As a practical use case, we select a dataset-specific perturbation level *λ*^∗^ as the largest noise level that retains a target fraction of duplicate pairings (e.g., ≥ 90%), and report split frequencies across replicate trees generated at *λ*^∗^ as support values. We validate the framework in a geometric toy model where evolving two-dimensional shapes provide a known ground-truth topology, and in an empirical globin benchmark using distances derived from 1 – TM-score. Across both settings, duplicate monophyly tracks the erosion of tree structure under distance-matrix noise, establishing an internally calibrated “resolution limit” for assigning confidence in distance-based structural phylogenetics.

## Introduction

Transformer-based models have become a foundation for modern structural biology [1], enabling structure prediction methods such as AlphaFold [2] and ESMFold [3]. This rapid expansion of the protein structure universe has created new opportunities for evolutionary analysis beyond sequence comparisons. In particular, structure-based similarity can retain signal in regimes where sequence identity is low, enabling the exploration of deep homology using three-dimensional folds [4, 5]. Tools such as Foldseek [6] allow rapid computation of normalised pairwise structural similarity scores (e.g., TM-score [7]), which can be converted to distance matrices (e.g., *d*_*ij*_ = 1−TM(*i, j*)) for neighbour-joining (NJ) tree construction [8, 9, 10].

A critical gap in this workflow is the estimation of statistical confidence. In classical sequence phylogenetics, the non-parametric bootstrap [11] is widely used: alignment columns are resampled to generate replicate datasets and a distribution of trees. Although the under-lying independence assumptions are biophysically imperfect due to epistasis and structural constraints [12], bootstrapping remains a practical convention for variance estimation in large-scale analyses.

For structure-derived distance matrices, however, there is no natural analogue of re-sampling alignment columns. A pairwise TM-score is a single scalar that summarises a global geometric superposition; it does not decompose into a set of discrete, approximately independent observations that can be resampled. Previous work has shown that resampling from molecular dynamics (MD)-generated conformational ensembles can provide a rigorous, physically grounded bootstrap analogue for structural phylogenies, by propagating thermo-dynamic (ensemble) uncertainty into replicate distance matrices and trees [8, 13]. However, although this ensemble-based resampling is the most principled route to confidence estimation, it remains computationally prohibitive for high-throughput settings involving many taxa or in web-based settings e.g., programs in the Structome suite [9, 10]. Prior work has shown that conformational ensembles can induce variability in pairwise structural similarity scores: for example, Malik et al. [13] compared fixed reference frames to full MD trajectories and observed pairwise score fluctuations. In the present work, we do not attempt to reproduce ensemble-resampled bootstraps directly; instead, we treat distance-space perturbations as a calibrated surrogate for such ensemble-induced variation, with calibration provided internally by the Duplicate Monophyly Criterion.

A computationally feasible alternative is *parametric bootstrapping* [14, 15], in which the distance matrix is perturbed according to a theoretical model and trees are reconstructed from replicate matrices. The key limitation is calibration: the variance parameter (*σ*^2^) is rarely known, and support values can become a tunable artefact of noise strength. Too little perturbation yields spuriously high support, whereas too much perturbation yields essentially random trees.

To address this calibration problem, we propose an empirical, data-driven strategy: the *Duplicate Monophyly Criterion* (DMC). The core intuition is self-consistency: if a perturbation regime is strong enough that the trivial relationship between a structure and its exact duplicate is no longer recovered, then that regime has overwhelmed subtler evolutionary signal. We operationalise this by expanding the distance matrix to include a virtual duplicate for every taxon, assigning each original–duplicate pair a small baseline distance, and tracking *duplicate monophyly D*(*λ*) (the “duplicate cherry survival” rate), i.e. the fraction of duplicate pairs recovered as exclusive two-tip clades. The breakdown of this signal defines an empirical “resolution limit”: the maximum perturbation level under which identity-level relationships remain reliable.

We develop this framework in two stages. First, we put together a geometric toy model in which two-dimensional polygons are made to evolve along a known bifurcating tree. Pair-wise Procrustes distances are computed between these simulated shapes and the resulting distance matrix is perturbed using a heteroscedastic, floor-augmented noise model, allowing us to study how noise, topological recovery, and duplicate cherry survival co-vary when ground truth is known. In the second stage, we use real protein structures by applying the same distance-space perturbation model to a distance matrix derived from 1 − TM-score. In both these settings, duplicates act as internal gauges (controls) of reliability, providing a principled calibration of perturbation strength and enabling bootstrap-like support estimation for distance-based structural phylogenetics. As a practical use case, we use the DMC to select a dataset-specific perturbation level *λ*^∗^ (the largest noise level that retains a target fraction of duplicate *cherries*), and we then summarise the robustness of the inferred topology by reporting *split* frequencies across replicate trees generated at *λ*^∗^.

## Mathematical Framework

To apply parametric bootstrapping to distance-based phylogenetics, we define a noise model that reflects two sources of uncertainty: the *heteroscedastic* error inherent in structural divergence (where uncertainty scales with distance), and the *systematic* error floor inherent in alignment or measurement limits (where some uncertainty exists even for very similar objects).

Let **D** be an *N* × *N* matrix of observed pairwise distances between objects, with entries *d*_*ij*_. We view these observed distances as noisy realisations of unknown “true” evolutionary distances *δ*_*ij*_:

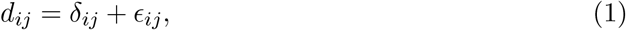

where *ϵ*_*ij*_ is a random perturbation term. We model *ϵ*_*ij*_ as a Gaussian random variable centred at zero. In contrast to simple multiplicative models where the variance vanishes as *d*_*ij*_ → 0, we introduce a floor-augmented heteroscedastic model:

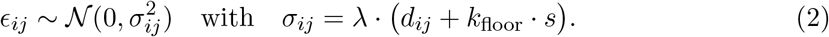

Here, *λ* is the global *noise level* (the parameter swept during calibration), *k*_floor_ is a fixed scaling constant (set to 2.5 in our experiments), and *s* is a dataset-specific scale factor defined as the median of all positive off-diagonal distances in the matrix being perturbed.

The term *k*_floor_ · *s* ensures that a baseline level of variance is applied to all edges, pre-venting very small distances from being artificially immune to perturbation.

In practice, we generate a perturbed distance matrix 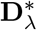 by drawing

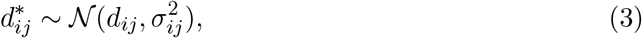

sampling once for *i* < *j* and enforcing symmetry 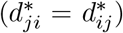, setting 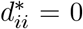, and clipping sampled distances to the bounded range [0, 1] after perturbation.

Given a perturbed matrix 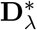, we reconstruct a neighbour-joining tree [16] and compare its topology to a reference tree obtained from the unperturbed matrix **D**. Because neighbour-joining produces unrooted trees, we quantify *topological accuracy A*(*λ*) using the retention of unrooted splits (bipartitions) [17]. Let 𝒮_ref_ denote the set of non-trivial bipartitions in the reference tree, and 𝒮_*λ*_ the corresponding set for the replicate tree. The accuracy is defined as

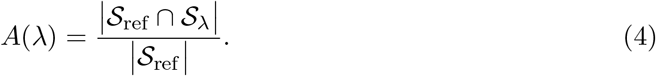

### The Duplicate Monophyly Criterion

The central difficulty in parametric bootstrapping is the choice of the noise level *λ*. Without an empirical way to constrain *λ*, support values can be arbitrarily tuned. To address this, we introduce an internal calibration mechanism based on synthetic taxon duplicates.

We construct an augmented dataset of size 2*N* by introducing, for each taxon *S*_*i*_, a virtual duplicate *S*_*i*_′ . At the level of the distance matrix, we form a 2*N* × 2*N* block matrix

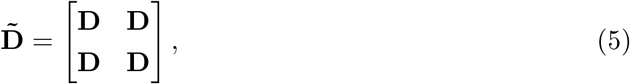

and relabel the taxa so that the first *N* entries correspond to {*S*_1_, …, *S*_*N*_} and the next *N* entries to {*S*_1_′, …, *S*_*N*_′}.

We then overwrite the cross-block diagonal entries 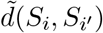 and 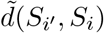 for all *i* with the non-zero tripwire distance defined in Eq. (6).

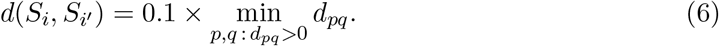

This value is one order of magnitude smaller than the finest non-zero resolution already present in the dataset, placing duplicates at a strictly finer distance scale than any observed non-identical pair and making duplicate pairing sensitive to the injected perturbation.

We apply the floor-augmented noise model in Eq. (2) to this expanded matrix. For a given *λ*, we generate a perturbed matrix 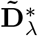 and reconstruct a neighbour-joining tree. For each pair (*S*_*i*_, *S*_*i*_′), we check whether they form a *two-tip cherry*, operationalised as follows: we compute the most recent common ancestor (MRCA) of *S*_*i*_ and *S*_*i*_′ in the NJ tree, and test whether the set of tips subtended by this MRCA is exactly {*S*_*i*_, *S*_*i*_′}. We define the *duplicate monophyly D*(*λ*) as the fraction of duplicate pairs satisfying this condition:

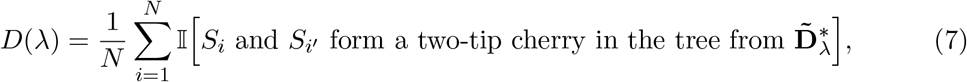

where 𝕀[·] is the indicator function.

### Resolution Limit and Calibrated Bootstrapping

The Duplicate Monophyly Criterion postulates that *D*(*λ*) provides a conservative, internal check on perturbation strength. If the injected noise is sufficient to disrupt the trivial grouping of duplicates (despite their minimal tripwire distance in Eq. (6)), then the perturbation regime has exceeded the local resolution of the dataset and is unlikely to preserve more subtle phylogenetic signal.

We define an empirical *resolution limit* by choosing a threshold *τ* ∈ (0, 1) (e.g. *τ* = 0.9) and identifying the largest noise level *λ*_*τ*_ for which duplicate monophyly (as measured by *D*(*λ*)) remains above this threshold:

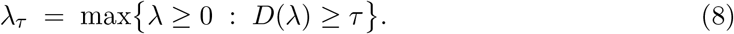

This *λ*_*τ*_ provides an empirical upper bound on the perturbation regime under which identity-level relationships remain stable. In practice, *λ* is evaluated on a finite grid, and *D*(*λ*) is estimated as the mean duplicate monophyly score across replicate perturbations at each grid point; we then set *λ*_*τ*_ to the largest tested *λ* for which the estimated *D*(*λ*) ≥ *τ* .

We then fix *λ* = *λ*_*τ*_ and generate a collection of perturbed *duplicated* distance matrices

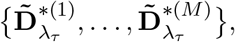

reconstruct a neighbour-joining tree from each replicate, and compute split support values on the original taxa by pruning duplicate tips and tallying the frequency of reference splits across replicates. These split frequencies provide bootstrap-like support values for the distance-based phylogeny. Importantly, the resulting support values reflect robustness to DMC-calibrated distance perturbations, intended as a computationally efficient surrogate for ensemble-based (MD/MC) resampling of structural conformations.

In the following sections, we instantiate this framework in two settings: first, a geometric toy model in which two-dimensional shapes evolve on a known tree and Procrustes distances are perturbed in matrix space (Realisation 1), and second, an empirical protein structural dataset in which distances are derived from 1 − TM-score and perturbed using the same model (Realisation 2).

### Realisation 1: Geometric Shapes and Distance-Matrix Noise

In the first realisation, we consider an idealised system where each taxon is a planar shape evolving along a known binary tree. We initialise the system with a regular polygon on the unit circle (with *n* = 20 vertices). This ancestral shape is propagated along a full binary tree of depth *G* = 4. At each branch, we generate descendant shapes by adding small isotropic Gaussian perturbations to the vertex coordinates,

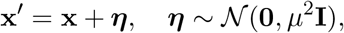

followed by recentering on the origin. We fix the mutation rate at an intermediate value (*µ* = 0.05) to generate a representative dataset where the tip shapes exhibit moderate geometric divergence but distinct clade structure. Additional mutation rates were explored, and all showed the same qualitative trend; we report results for *µ* = 0.05.

Distances between tip shapes are defined using Procrustes analysis [18]: for each pair (*i, j*), we compute the Procrustes distance *d*_*ij*_ and assemble these into the baseline distance matrix **D**. To match a bounded structural distance scale, we normalise **D** by its maximum entry so that *d*_*ij*_ ∈ [0, 1]. We then apply the floor-augmented matrix-based noise model in Eq. (2) to this fixed **D**, construct duplicate-augmented matrices as in Eqs. (5)–(6), and evaluate both *A*(*λ*) and *D*(*λ*) (Eqs. (4) and (7)) across a sweep of *λ*. Because the underlying tree is known by construction, this setting provides a controlled benchmark for assessing whether distance-matrix perturbations diagnose the loss of topological signal induced by accumulated geometric perturbations.

### Realisation 2: Protein Structures and Distance-Matrix Noise

In the second realisation, we apply the framework to real protein structures, where explicit coordinate-space simulation (e.g. MD) is computationally prohibitive. Here, we operate entirely in distance space.

We use a benchmark of globin structures (including *α*-haemoglobin [PDB accession IDs: 1hv4 A, 4hhb A, 1t1n A], *β*-haemoglobin [PDB accession IDs: 4hhb B, 1hv4 B, 1t1n B] and myoglobin [PDB accession IDs: 1myf A, 1myg A]). Pairwise structural similarity scores are computed using Foldseek and converted to distances as

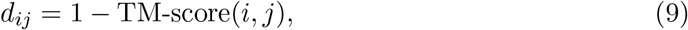

yielding a distance matrix **D** over the selected structures. We then construct the augmented matrix with duplicates separated by the tripwire distance in Eq. (6) and sweep the noise parameter *λ* using the floor-augmented model in Eq. (2). By tracking the decay of duplicate monophyly *D*(*λ*) (Eq. (7); defined via two-tip cherry recovery) alongside the stability of the reference neighbour-joining topology inferred from the unperturbed matrix, *A*(*λ*) (Eq. (4)), we identify a resolution limit *λ*^∗^ := *λ*_*τ*_ (Eq. (8)) at which bootstrap-like support values are computed for the inferred globin tree.

## Results

### Geometric toy model: evolution in shape space

We first asked whether a simple geometric system could provide a controlled benchmark in which the effects of distance-space noise on tree topology could be examined explicitly. To this end, we represented each taxon as a planar polygon with *n* = 20 vertices placed on the unit circle, and evolved this ancestral shape along a full binary tree of depth *G* = 4 (16 tips). At each branch, the parent polygon was perturbed by adding isotropic Gaussian noise to its vertex coordinates, followed by recentering on the origin. The branch-wise mutation rate *µ* controls the magnitude of these coordinate displacements and therefore the rate at which descendant shapes diverge from the ancestral circle.

Figure 1 illustrates the effect of two contrasting mutation regimes. At a low mutation rate (*µ* = 0.01), tip shapes remain visually close to the circle, and gradual deformation along the tree is visible in the internal nodes (Fig. 1, left). The overall geometry is well preserved and the tips differ only by small perturbations. In contrast, at a higher mutation rate (*µ* = 0.1) the polygons become progressively more jagged with depth, and the tip shapes are highly heterogeneous (Fig. 1, right). This simple system therefore captures the intuitive notion that stronger drift in shape space erodes the geometric signal supporting a well-resolved tree, while retaining a known underlying topology defined by the branching process.

**Figure 1:**
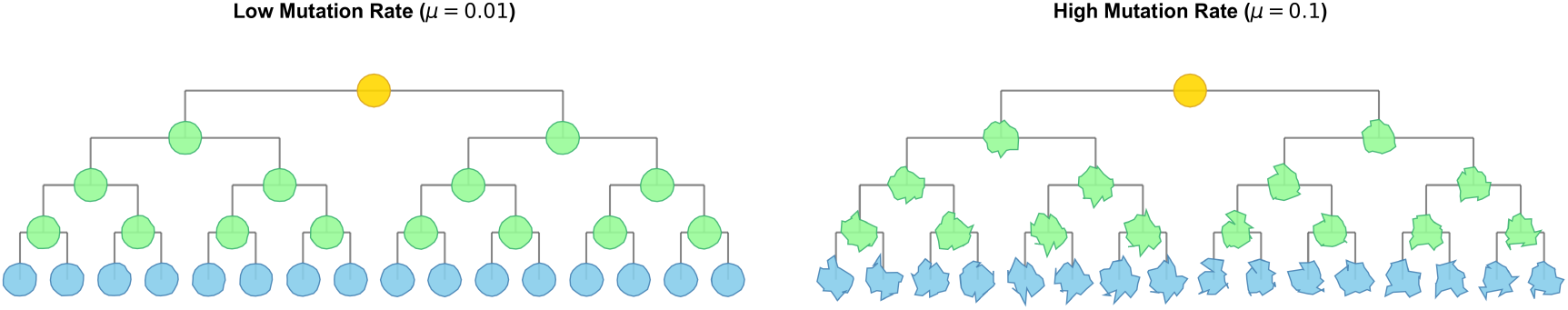
Visualisation of the geometric toy model. A regular 20-sided polygon (the ancestor) is propagated along a binary tree of depth *G* = 4 by adding Gaussian perturbations to vertex coordinates at each branch (Brownian increments in shape space), followed by recentering. **(Left)** At a low mutation rate (*µ* = 0.01), tip shapes retain the general circular form of the ancestor, qualitatively resembling conserved folds. **(Right)** At a high mutation rate (*µ* = 0.1), accumulated perturbations yield highly heterogeneous tip shapes, qualitatively resembling divergent structures with low similarity.

### Duplicate monophyly tracks topological accuracy in the toy system

We used this toy model to test whether DMC-style duplicate monophyly provides a useful proxy for the topological stability of a distance-based tree under distance-matrix perturbations. For these experiments, we fixed an intermediate mutation rate (*µ* = 0.05) to generate a single set of tip shapes and a corresponding “true” tree given by the known binary branching structure. Pairwise Procrustes distances between tip polygons were assembled into a baseline distance matrix **D**. The ground-truth set of unrooted splits, 𝒮_ref_, was defined directly from the planted branching process and used as the reference against which all subsequent neighbour-joining trees were compared.

To probe the effect of measurement noise, we applied the floor-augmented heteroscedastic noise model directly to the distance matrix, mirroring the intended procedure for real protein structures. For each value of the noise level *λ* in a grid from 0 to 0.5, we generated replicate matrices by adding noise defined by Eq. (2) and reconstructed neighbour-joining trees. Comparing the unrooted splits of these trees to 𝒮_ref_ yields the topological accuracy *A*(*λ*) (Eq. (4)). In parallel, we constructed an augmented distance matrix in which every tip shape was duplicated and separated by the “tripwire” distance (Eq. (6)). We applied the same matrix-space noise model to this expanded dataset and quantified the fraction of duplicate pairs that formed monophyletic cherries (two-tip cherries), giving the duplicate monophyly curve *D*(*λ*) (Eq. (7)).

The resulting curves are shown in Figure 2. At low noise (*λ* ≈ 0), both *A*(*λ*) and *D*(*λ*) are high, indicating that both the planted topology and the duplicate pairs are recovered. As *λ* increases, both quantities decline, but they do so at different rates: the topological accuracy (blue curve) decays more rapidly than the duplicate monophyly (red curve). This separation follows from the heteroscedastic noise model, in which larger pairwise distances receive larger absolute perturbations, whereas duplicate pairs are separated by a very small tripwire distance and are perturbed primarily by the noise floor. Consequently, duplicate monophyly provides an *internal gauge* of reliability across noise levels.

**Figure 2:**
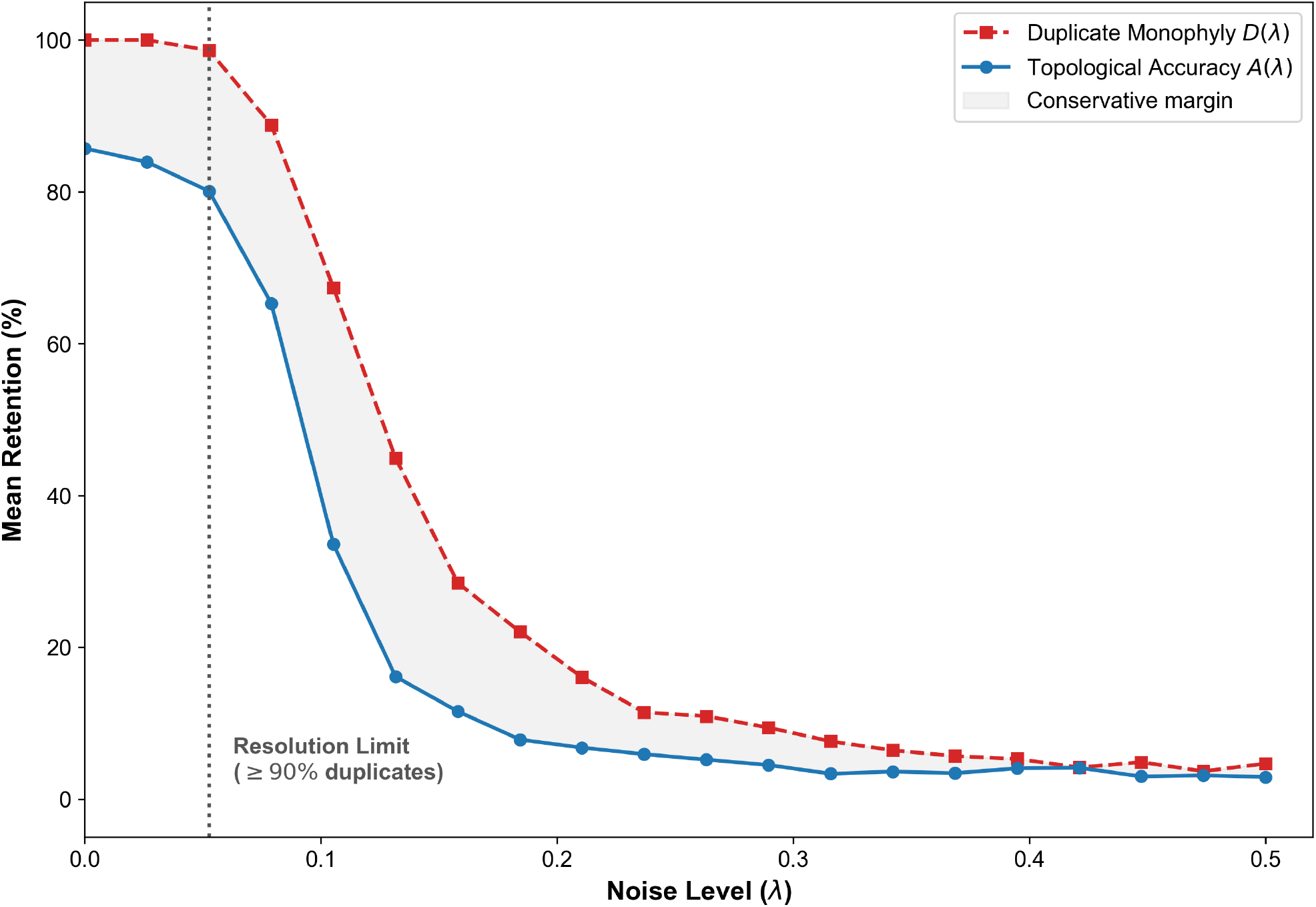
Duplicate monophyly and topological accuracy under distance-matrix noise in the geometric toy model. For a fixed geometric regime (mutation rate *µ* = 0.05), we applied the floor-augmented heteroscedastic noise model (Eq. (2)) to the Procrustes distance matrix and reconstructed neighbour-joining trees across a sweep of noise levels *λ* ∈ [0, 0.5]. The blue curve shows the mean topological accuracy *A*(*λ*), defined as the fraction of reference unrooted splits retained (Eq. (4)). The red dashed curve shows the duplicate monophyly *D*(*λ*), defined as the fraction of taxon–duplicate pairs recovered as cherries in the duplicate-augmented trees (Eq. (7)). The grey shaded region highlights the gap between *D*(*λ*) and *A*(*λ*), capturing the regime where duplicates still survive but deeper splits have begun to fail. The vertical dotted line marks the empirical resolution limit, defined as the largest *λ* for which *D*(*λ*) ≥ 90%. Beyond this point, both curves collapse, indicating that noise has overwhelmed the underlying topological signal.

We defined an empirical resolution limit as the largest noise level at which duplicate monophyly remains at or above 90%. This threshold (vertical dotted line in Figure 2) identifies a regime where the planted topology remains largely recoverable (here, *A*(*λ*) ≈ 80%). Beyond this point, as duplicate survival begins to fail, the topological signal has already degraded substantially, supporting the view that high duplicate monophyly is a necessary condition for interpreting deeper structure in the inferred tree.

### Globin structural phylogeny: parametric noise in distance space

We next asked whether the Duplicate Monophyly Criterion behaves similarly on real protein structural data, where distances arise from three-dimensional folds rather than from an idealised geometric system. To this end, we assembled a small benchmark of globin structures comprising representatives of *α*-haemoglobin, *β*-haemoglobin and myoglobin. Pairwise structural similarity scores were computed using Foldseek and converted to distances as

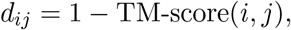

yielding an *N* × *N* distance matrix **D** over the chosen globin set (*N* = 8).

A neighbour-joining tree constructed from **D** served as the reference distance-based phylogeny, recovering the expected separation of myoglobins from the *α/β* haemoglobin clade. Because globin relationships are well established, we treat the canonical partitioning (myoglobin vs haemoglobin, and *α* vs *β* haemoglobin) as an empirical benchmark, and note that the unperturbed NJ tree recovered these expected splits.

In this empirical setting, we implemented the noise model directly on the duplicate-augmented dataset. First, we constructed an expanded 2*N* × 2*N* matrix by duplicating every taxon. Duplicate pairs (*S*_*i*_, *S*_*i*_′) were assigned a small but non-zero “tripwire” distance (0.1*×* the minimum non-zero distance in **D**), so that duplicates represent a finite resolution challenge on the dataset’s native scale rather than an exact identity.

We then applied the *floor-augmented heteroscedastic noise model* to this expanded matrix. For each value of the noise level *λ* on a regular grid between 0 and 0.5, we generated noisy realisations 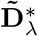 of size 2*N* × 2*N* by sampling

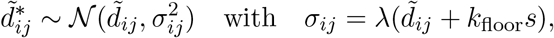

for all *i* ≠ *j*, enforcing symmetry, setting diagonal terms to zero, and clipping distances to the range [0, 1].

For each noisy realisation, we reconstructed a neighbour-joining tree and computed two metrics from the same tree:

1. **Duplicate Monophyly** *D*(*λ*): The fraction of duplicate pairs (*S*_*i*_, *S*_*i*_′) recovered as monophyletic two-tip cherries.
2. **Topological Accuracy** *A*(*λ*) **(reference split retention):** The tree was pruned to remove the duplicate tips {*S*_1_′, …, *S*_*N*_′}, and the unrooted splits of the remaining tree were compared to the reference globin tree (the unperturbed NJ tree) to calculate the fraction of reference splits retained.

This procedure ensures that both metrics reflect the resolution limit of the exact same perturbed data. Both *A*(*λ*) and *D*(*λ*) were estimated by averaging over 100 independent replicates for each value of *λ*.

The behaviour of *A*(*λ*) and *D*(*λ*) for the globin dataset is summarised in Figure 3. At *λ* = 0, both curves are at 100 % (as expected, because the reference splits are defined from the neighbour-joining tree on the unperturbed matrix), and each taxon pairs cleanly with its duplicate. As the noise level increases, both *A*(*λ*) and *D*(*λ*) decline, with duplicate monophyly consistently slightly lower than reference split retention. This indicates that duplicate behaviour closely tracks the erosion of the reference topology and can be used as an empirical gauge of how much parametric noise the dataset can tolerate before the reference relationships become unreliable.

**Figure 3:**
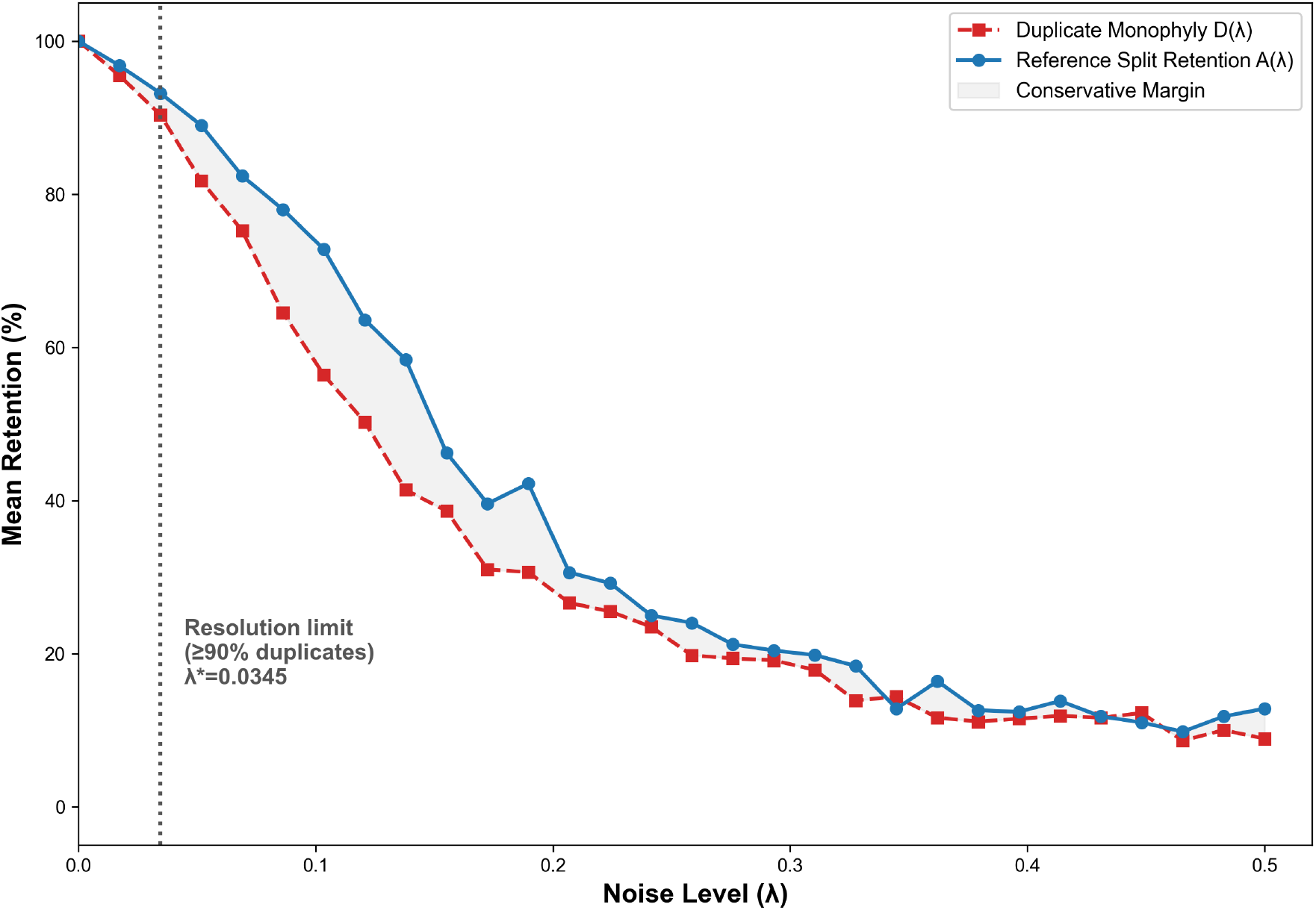
Calibration of parametric bootstrapping for globin structures. Results for the globin dataset (*N* = 8, 100 replicates per noise level). The blue curve shows reference split retention *A*(*λ*) calculated after pruning duplicate tips. The red dashed curve shows duplicate monophyly *D*(*λ*) calculated on the full tree. Both metrics are derived from the same set of noise-perturbed 2*N* × 2*N* matrices. In this dataset, *D*(*λ*) lies slightly below *A*(*λ*), indicating that the tripwire pairing provides a strict local-resolution test relative to the reference split set. The vertical dotted line marks the empirical resolution limit (largest *λ* with *D*(*λ*) ≥ 90%; here *λ*^∗^ ≈ 0.0345), identifying a conservative perturbation regime for support estimation.

We defined an empirical resolution limit as the largest noise level at which duplicate monophyly remains at or above 90 %. For the globin dataset this occurs at *λ*^∗^ ≈ 0.0345, at which point reference split retention remains high (*A*(*λ*^∗^) ≳ 90%). We then fixed *λ* = *λ*^∗^ and generated 100 replicate trees to estimate support for the reference splits. This yielded 100% support for the major splits separating myoglobins, *α*-globins and *β*-globins, while internal structure within the haemoglobin subclades showed more variable support (e.g. 96% for one internal *α*-globin split and 65% for an internal *β*-globin split). These results support the use of *D*(*λ*) as a conservative, empirically accessible gauge for calibrating parametric perturbation strength in distance-based structural phylogenies.

## Discussion

The rapid accumulation of protein structural data has necessitated a shift from sequence-based to structure-based phylogenetics. However, this transition introduces a fundamental challenge: the loss of the standard non-parametric bootstrap. In sequence analysis, confidence is estimated by resampling alignment columns, a method predicated on the assumption of independent, discrete characters [11]. Structural comparisons, summarised by a continuous metric like TM-score, lack these discrete units, resulting in the absence of a computationally tractable native framework for statistical validation. While parametric bootstrapping offers a theoretical alternative, it requires an input noise parameter that is typically unknown. Without a calibrated way to determine how much variance to inject, support values can be arbitrarily inflated or deflated.

To address this calibration problem, we proposed the Duplicate Monophyly Criterion as an internal, empirical strategy. The logic is grounded in a necessary-condition argument: while the true deep topology of an empirical tree is unknown, the relationship between a taxon and its exact duplicate is known *a priori* to be monophyletic. We hypothesised that the stability of these synthetic “cherries” could serve as a proxy for the stability of the tree as a whole. Specifically, a perturbation regime that disrupts the grouping of duplicates indicates that the signal-to-noise ratio has degraded substantially. Therefore, any practical parametric bootstrap intended to reflect meaningful uncertainty should operate in a regime where these control relationships are largely preserved.

We first evaluated this conjecture in a controlled geometric toy model (Fig. 1), where shapes evolve along a known binary tree. In this setting, the underlying topology is known by construction, allowing the retention of planted unrooted splits to be measured directly. When we applied the floor-augmented heteroscedastic perturbation model in *distance space* (Fig. 2), the decay of duplicate monophyly tracked the decay of split retention as noise increased. Moreover, duplicate monophyly decayed slightly more slowly than split retention, consistent with the interpretation that the duplicate criterion provides a conservative necessary condition: once duplicates fail, the planted topology has already degraded substantially.

We then extended the framework to empirical protein structural data. Ideally, structural uncertainty would be estimated by explicit simulation, where alternative conformations extracted from Molecular Dynamics (MD) trajectories represent coordinate-space variance [8, 13]. While biologically rigorous, generating conformational ensembles for every taxon in a large phylogeny is computationally intractable. To achieve scalability, we instead approximate such fluctuations by applying a floor-augmented heteroscedastic noise model directly to the (duplicate-augmented) distance matrix. This allows for the rapid generation of many replicate trees without the need for expensive physical simulations.

A practical feature of this implementation is the definition of a finite duplicate separation (the “tripwire” distance). Rather than treating duplicates as exact identities, we assign each original–duplicate pair a small but non-zero distance set to one order of magnitude below the smallest non-zero distance already present in the dataset. This ensures that duplicates represent a strict local-resolution test on the dataset’s own distance scale, rather than an idealised case of perfect identity. In the globin benchmark (Fig. 3), duplicate monophyly decayed slightly *faster* than reference split retention, indicating that the tripwire pairing can function as a conservative gauge of stability under matrix perturbation. In practice, calibrating the perturbation level to retain a target fraction of duplicates therefore yields a cautious operating regime for support estimation.

### Disclaimer

It is important to acknowledge that, like all heuristic approaches in computational biology, the Duplicate Monophyly Criterion represents a strategic compromise between computational efficiency and physical realism. We note that the primary intended deployment context for the DMC is web-based structural phylogenetics tools operating on datasets of tens to low hundreds of structures—such as those in the Structome suite—where MD-based ensemble generation is computationally infeasible and a scalable, internally calibrated confidence measure is most needed. In principle, the most faithful way to estimate structural uncertainty is to generate conformational ensembles through MD or Monte Carlo simulations, which directly sample the underlying thermodynamic landscape [8, 13]. However, such simulations remain prohibitively expensive for large-scale analyses. By approximating these fluctuations directly in distance space, our framework enables statistical validation in settings where rigorous biophysical simulation is infeasible. While the inclusion of duplicates increases the dimensionality of the neighbour-joining inference (*N* → 2*N*), the tripwire distance is chosen so that each taxon–duplicate pair is strongly favoured to form a two-tip cherry under neighbour-joining. Because neighbour-joining is global and auxiliary taxa can in principle influence intermediate joins, support values are computed after pruning duplicate tips and are interpreted as stability of the original taxa under DMC-calibrated perturbations.

This comes at the cost of sacrificing the granular accuracy of explicit ensemble sampling, but it provides a scalable and internally calibrated mechanism for assessing the robustness of distance-based structural phylogenies. We emphasise that the perturbation model is an empirical surrogate: it does not explicitly model coordinate-space dynamics, non-Gaussian conformational heterogeneity, or correlations among distances induced by shared structural features. The floor term and clipping to [0, 1] are practical stabilisers rather than biophysical statements, and the resulting support values should be interpreted as stability under calibrated distance perturbations rather than as a literal sampling probability from an explicit structural generative model. The Duplicate Monophyly Criterion should therefore be viewed not as a replacement for MD-based uncertainty quantification, but as a practical alternative that allows confidence estimation to remain tractable at scale.

Together, these results position the Duplicate Monophyly Criterion as a practical and principled foundation for bringing empirically calibrated support values to distance-based structural phylogenetics.

## Conclusion

Distance-based structural phylogenies lack a computationally tractable native framework for statistical validation. In this work, we introduced the Duplicate Monophyly Criterion, a calibration strategy that uses synthetic taxon duplicates to empirically determine the perturbation tolerance of a dataset. Across both a geometric toy model and an empirical globin benchmark, we showed that the stability of duplicate pairings closely tracks the retention of tree structure under distance-matrix noise, providing an internal gauge for selecting a conservative perturbation regime. This yields a practical workflow: researchers can identify a “resolution limit” where duplicate monophyly remains high (e.g. ≥ 90%) and use the corresponding noise level to calibrate parametric bootstrapping and compute split frequencies as support values. By providing a scalable alternative to explicit Molecular Dynamics ensemble generation, this framework enables calibrated confidence estimation for distance-based structural phylogenies, supporting the interpretation of inferred trees as testable evolutionary hypotheses. As a practical use case, this calibration step can be integrated into existing distance-based structural phylogeny web resources (e.g. tools that currently return a single neighbour-joining tree such as Structome-TM), enabling support values to be reported alongside the inferred topology without requiring MD/MC simulation infrastructure.

## Availability and implementation

The Duplicate Monophyly Criterion has been implemented as a new module within Structome Playground, an interactive educational web resource available at https://biosig.lab.uq.edu.au/structome_playground/. While the platform’s initial modules (1–3) allow users to explore the fundamental relationships between geometric shape, distance matrices, and neighbour-joining topology, the newly added Module 4: The Resolution Limit, explicitly implements the calibration framework described in this work. This module allows users to simulate the effect of noise in real-time, visualising how the stability of duplicate pairs (Duplicate Monophyly; two-tip cherry recovery) functions as a conservative necessary condition for topological accuracy under increasing matrix noise. The server provides an accessible, no-code environment for verifying the logic of the method and building intuition for distance-based calibration.

## Funding

DBA is supported by the investigator grant from the National Health and Medical Research Council (NHMRC) of Australia [GNT1174405] and the Victorian government Operational Infrastructure Support program.

## Generative AI statement

The authors declare that Gen AI was used in the creation of this manuscript.

## Notes

### Competing Interest Statement

The authors have declared no competing interest.

